# Architecture of the human G-protein-methylmalonyl-CoA mutase nanoassembly for B_12_ delivery and repair

**DOI:** 10.1101/2023.03.23.533963

**Authors:** Romila Mascarenhas, Markus Ruetz, Harsha Gouda, Natalie Heitman, Madeline Yaw, Ruma Banerjee

## Abstract

G-proteins function as molecular switches to power cofactor translocation and confer fidelity in metal trafficking. MMAA, a G-protein motor, together with MMAB, an adenosyltransferase, orchestrate cofactor delivery and repair of B_12_-dependent human methylmalonyl-CoA mutase (MMUT). The mechanism by which the motor assembles and moves a >1300 Da cargo, or fails in disease, are poorly understood. Herein, we report the crystal structure of the human MMUT-MMAA nanomotor assembly, which reveals a dramatic 180° rotation of the B_12_ domain, exposing it to solvent. The nanomotor complex, stabilized by MMAA wedging between two MMUT domains, leads to ordering of the switch I and III loops, revealing the molecular basis of mutase-dependent GTPase activation. The structure explains the biochemical penalties incurred by methylmalonic aciduria-causing mutations that reside at the newly identified MMAA-MMUT interfaces.

## Introduction

Intracellular trafficking pathways shepherd rare, reactive, but essential metal and organic cofactors that undergird metabolism. With B_12_, clinical genetics studies (1) had provided early insights into the multitude of handlers that shepherd and repair this cofactor, which is needed by just two mammalian enzymes: cytoplasmic methionine synthase and mitochondrial methylmalonyl-CoA mutase (MMUT) (2,3). In both enzymes, the cobalt ion in B_12_ (or cobalamin) cycles between different oxidation states and is susceptible to inactivation (4). In methionine synthase, inactive cob(II)alamin is repaired *in situ* in the presence of an electron and a methyl group donor, to restore the active methylcobalamin form (5,6). In MMUT, the inactive cob(II)alamin is physically off-loaded onto adenosyltransferase (MMAB) in a process powered by GTP hydrolysis and catalyzed by a second chaperone, MMAA (Figure 1A) (7-9). MMAB converts cob(II)alamin to 5’-deoxyadenosylcobalamin (AdoCbl) in the presence of ATP and an electron donor, and then re-loads the active cofactor form onto MMUT (7,9,10). Structural insights into how such a gargantuan cofactor (1329 or 1579 Da for inactive/active forms) is loaded/off-loaded, and the inter-protein signals that orchestrate these processes, have been elusive. Mutations in each of the three mitochondrial B_12_ trafficking proteins are associated with hereditary methylmalonic aciduria, an inborn error of metabolism with a prevalence of ∼1:90,000 births (1). MMUT catalyzes the isomerization of methylmalonyl-CoA (M-CoA) funneled from the catabolic VOMIT (valine, odd-chain fatty acid, methionine, isoleucine and threonine) pathway reactions to succinyl-CoA, which enters metabolic mainstream (11). MMUT deficiency leads to the anaplerotic insufficiency of TCA cycle intermediates that can potentially be restored by a membrane soluble form of α-ketoglutarate (12).

**Figure 1.**
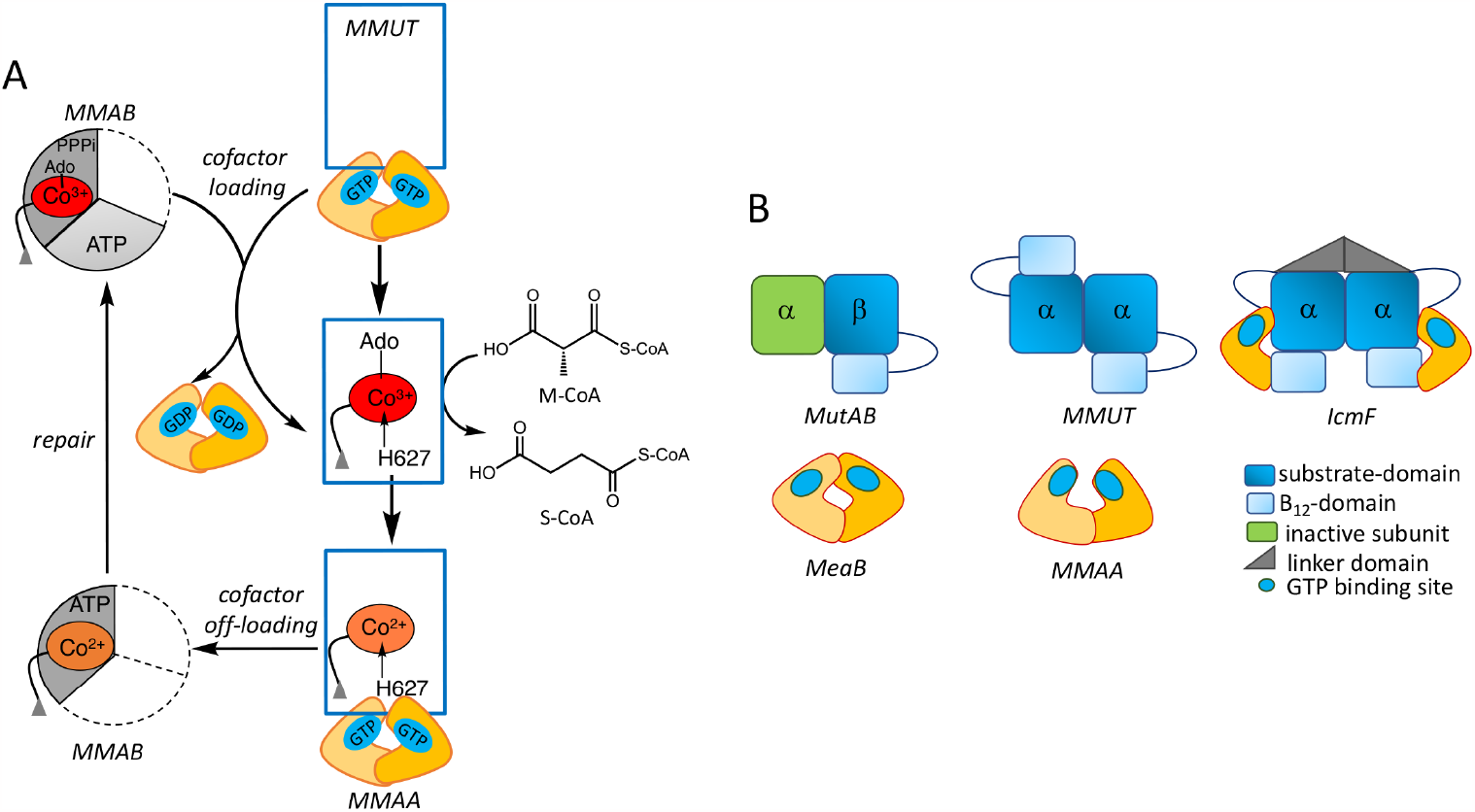
Mitochondrial human B_12_ trafficking proteins and comparison in their organization with bacterial homologs. **A**. Model for cofactor loading and off-loading in the mitochondrial B_12_- trafficking pathway. After AdoCbl is loaded onto MMUT (blue) by MMAB (grey), MMUT catalyzes the isomerization of methymalonyl-CoA (M-CoA) to succinyl-CoA (S-CoA). The GAP function of MMUT enhances the GTPase activity of MMAA (yellow), which assists in the transfer of cofactor to/from MMUT. MMAB is an adenosyltransferase and converts cob(II)alamin to AdoCbl and loads it onto MMUT. **B**. Cartoons showing the topological differences between bacterial MutAB, human MMUT and IcmF.

MMUT initiates substrate isomerization by homolytic cleavage of the cobalt-carbon bond in AdoCbl, generating cob(II)alamin and the 5’-deoxyadenosyl radical (13). Inadvertent loss of the 5’-deoxyadenosine moiety from the MMUT active site leads to inactive enzyme with cob(II)alamin bound (Figure 1A) (14). The structural basis for how apo- or inactive human MMUT signals to and recruits MMAA and MMAB for cofactor loading/off-loading is not known. Our understanding of how MMAA gates cofactor transfer to/from MMUT has emerged primarily from studies on the homologous proteins in *Methylobacterium extorquens (7,10,14-17)* and the *Cupriavidus metallidurans* fusion protein Icmf, comprising isobutyryl-CoA mutase and its G-protein chaperone(18,19). While the bacterial and human orthologs of MMAA and MMUT share ∼50% sequence identity, they exhibit large topological differences (Figure 1B), which limit their utility as models of the human proteins and the biochemical penalties associated with their disease-causing variants.

The *M. extorquens* orthologs of MMAA (MeaB) and MMUT (MutAB) are tightly bound; their affinity is modulated by the G-nucleotide and cobalamin ligands bound to the respective proteins (16,17). MeaB is a versatile chaperone that assists AdoCbl transfer from MMAB to MutAB, and cob(II)alamin transfer in the opposite direction, while also protecting the cofactor on MutAB against inactivation (14,16,17,20-22). The canonical G-protein switch I and II loops afford nucleotide-responsive allosteric regulation of MeaB functions and suppress its intrinsic GTPase activity (20,22). A third conformationally plastic switch III loop is important for the gating and editing functions of MeaB, which are corrupted by patient mutations that localize to this region (8,20). Crystal structures of MeaB have captured the switch III loop in multiple nucleotide-sensitive poses (19,20). Switch I and III loops are disordered in the structure of human MMAA, limiting insights (23).

Each IcmF monomer comprises an N-terminal B_12_-binding Rossmann fold domain, a nucleotide binding G-domain, a structured linker forming the dimer interface, and the C-terminal substrate-binding TIM barrel domain (Figure 1B) (18). While the G-domains in IcmF are structurally homologous to MeaB and MMAA, they are located at opposite ends, precluding dimer formation. The conserved nucleotide binding site is located at the interface between the substrate and G-domains.

MeaB and MMAA are homodimers with low intrinsic GTPase activity, which are activated in complex with the cognate mutase (17,24). The human and bacterial G-proteins have distinct interfaces and their nucleotide binding sites reside on the same or opposite faces of the dimer, respectively at distances of 25 Å (MMAA) and 50 Å (MeaB). The oligomeric composition of the mutase is different between the bacterial (αβ, MutAB) and human (α_2_) homologs, which translate into architectural differences in their complexes with the corresponding G-proteins (Figure 1B). The bacterial proteins exist in a 2:1 complex of MutAB:MeaB (22). A recent structure of the complex between the B_12_ domain of MutAB and MeaB in the presence of GMPPCP, captured a large conformational change, leading to the reorganization of the MeaB switch III loop and organization of the GTPase site (25). In contrast to the *M. extorquens* assembly, the human proteins exist as a variety of free and equilibrating oligomeric complexes varying from linear to annular with a predominant stoichiometry of 1:1 MMUT:MMAA (designated M_n_C_n_ where n = 2 as in the M_2_C_2_ complex (8). Patient mutations in the switch III region of MMAA impact the distribution between these oligomeric forms (8).

Herein, we report the crystal structure of the human M_2_C_2_ complex with coenzyme A (CoA) and GDP, which reveals dramatic conformational changes that occur as a prelude to cofactor translocation. MMAA stabilizes a 180° rotation of the B_12_ in relation to the substrate domain in MMUT by wedging between the two and undergoes ordering of its switch I and III loops. In this exploded conformation, the B_12_ domain moves from a sequestered to a solvent exposed pose, revealing how it is primed for cofactor loading/off-loading and explaining the mechanistic basis of pathogenic mutations that localize at the new interfaces formed in the M_2_C_2_ nanomotor complex.

## Results

The crystal structure of the human M_2_C_2_ complex with CoA and GDP•Mg^2+^ was solved by molecular replacement at a resolution of 2.8 Å (Table S1). Each molecule in the asymmetric unit contains two dimers each of MMUT and MMAA (Figure 2A), forming an annular structure as seen previously by negative staining (8). Each monomer in the MMUT α_2_ dimer comprises an N-terminal TIM barrel domain that binds substrate and a C-terminal Rossman fold domain that binds B_12_ (23). The substrate domains form the interface in the head-to-toe dimer arrangement, positioning the two B_12_ domains at opposite ends (Figure 2B). The substrate domain of an MMUT monomer forms an interface with one chain of MMAA, while the B_12_ domain interacts with a second MMAA chain (Figure 2C). Conversely, one chain in each MMAA α_2_ dimer simultaneously interacts with the substrate domain from one MMUT and the B_12_ domain from the second MMUT, leading to the annular form (Figure 2D).

**Figure 2.**
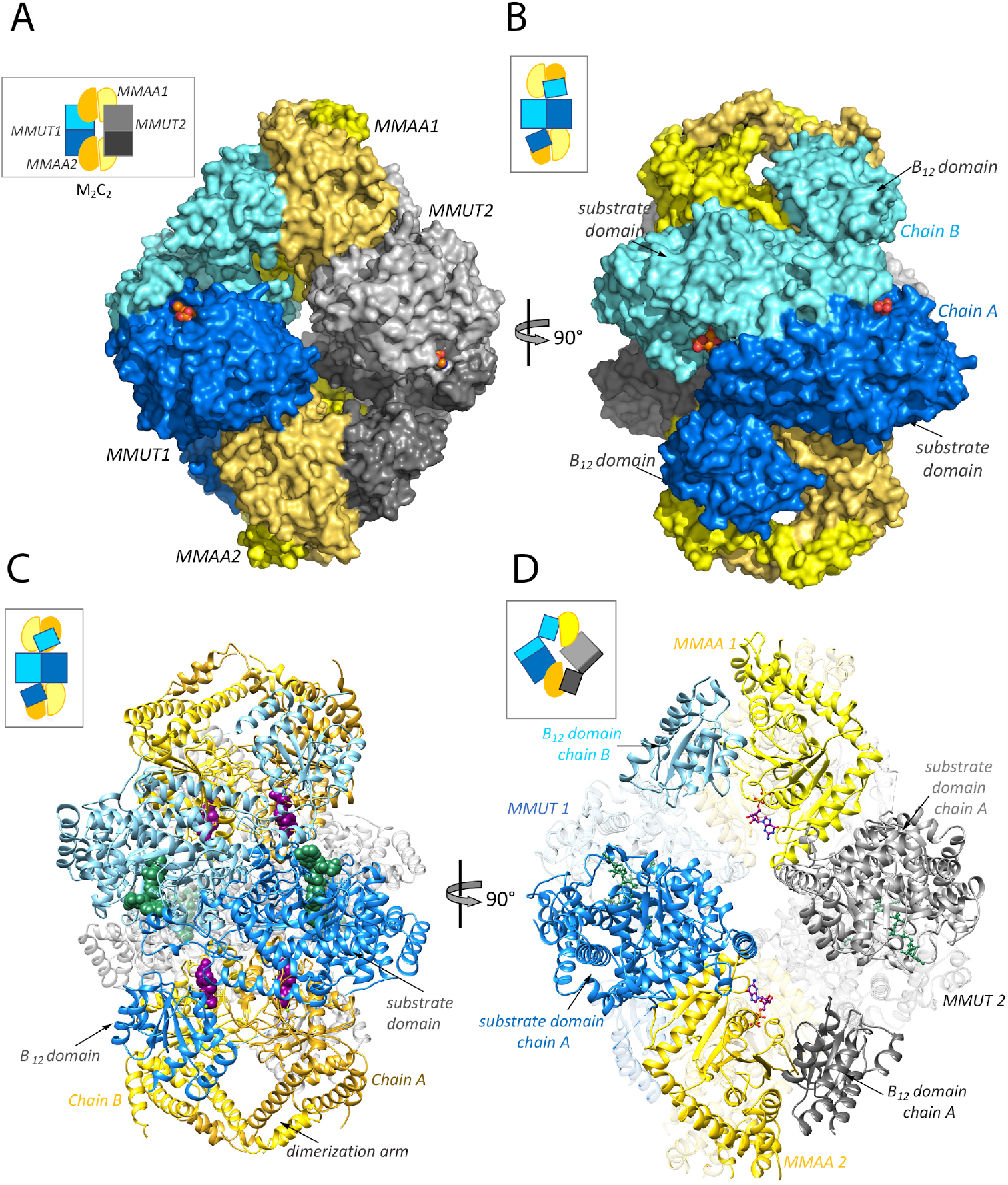
Structure of the human M_2_C_2_ complex. **A**. Surface representation of the M_2_C_2_ structure. MMUT1 and 2 are shown in blue and grey, respectively and the dark and light shades of MMAA (yellow) represent the monomeric chains in each α_2_ dimer. **B**. A 90° rotation shows that each MMUT chain (dark and light blue) interacts with one MMAA dimer. **C, D**. Ribbon representation of the M_2_C_2_ complex. GDP (purple) and CoA (50) are shown as spheres. **D**. One chain in MMAA2 (gold) simultaneously interacts with the substrate-domain in MMUT1 (dark blue) and the B_12_-domain in MMUT2 (dark grey). GDP is shown as purple sticks.

### MMUT in the M_2_C_2_ complex

The B_12_ is rotated ∼180° in the M_2_C_2_ complex versus in MMUT, which is evident from overlays of MMUT in the complex on apo-MMUT (PDB code 3BIC), MMUT•AdoCbl (PDB code 2XIJ) or MMUT•CoA•AdoCbl (PDB: 2XIQ) (Cα RMSD across all pairs=17 Å) (Figure 3A, Movie S1). MMAA is wedged between the two domains, stabilizing the exploded MMUT conformation (Figure S1A) that is secured by multiple interactions between them. The swinging out of the B_12_ domain is enabled by a flexible interconnecting belt (∼100 residues), which is partially unresolved in the M_2_C_2_ structure (between 580-595), suggesting mobility. In free MMUT, AdoCbl is nestled between the stacked B_12_ and substrate domains and shielded from solvent by an ordered belt (Figure 3A). In the M_2_C_2_ complex, this interface is disrupted by movement of the B_12_ domain, which exposes it to solvent. The MMUT loop (residues 622-629) that carries the conserved B_12_ ligand (His-627), is partially disordered in the “unstacked” conformation. The CoA threads through a channel in the substrate domain inducing it to close in on itself, as seen previously in the structure of MMUT bound to a substrate analog (23).

**Figure 3.**
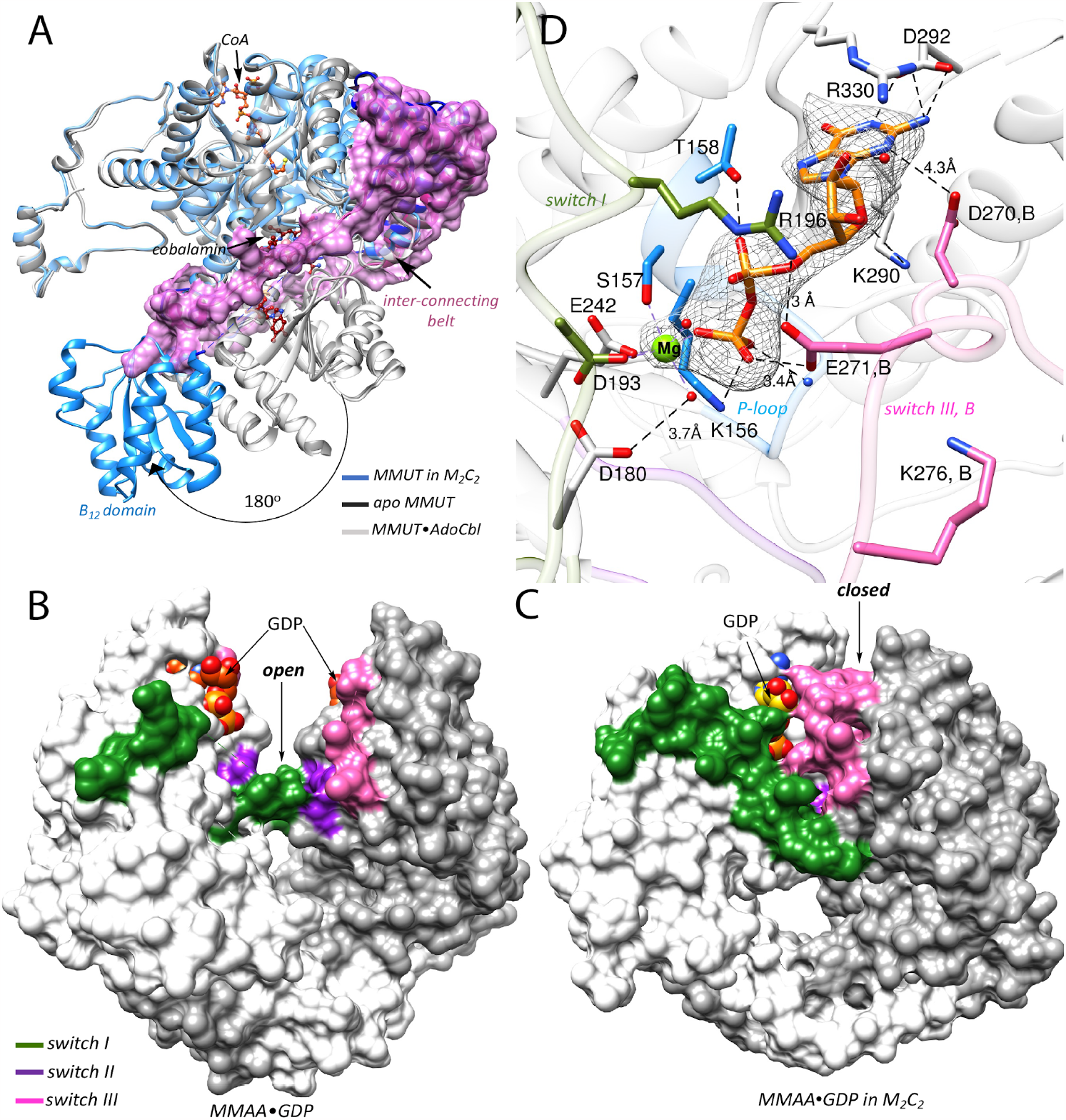
**A**. Overlay of MMUT in the M_2_C_2_ complex (blue) on free apo-MMUT (PDB: 3BIC) (dark grey) or with CoA (orange) and AdoCbl (maroon) bound (PDB:2XIQ) (light grey) shows that the B_12_ domain is swung out by ∼180° and is solvent exposed. In MMUT•AdoCbl an interconnecting belt (purple surface) wraps around the active site and protects the cofactor from solvent. In M_2_C_2_ the interconnecting belt is partially disordered (dark blue) **B**,**C**. Comparison of free MMAA (PDB: 2WWW) (B) and MMAA in M_2_C_2_ (C), shows that it undergoes a large shift from an open to a closed conformation in the M_2_C_2_ complex, which leads to ordering of the switch I (50) and III (pink) loops. **D**. The MMAA active site in M_2_C_2_ shows that GDP (orange sticks) binding is stabilized by multiple hydrogen bonds as described in the text. Asp-270 and Glu-271 from the switch III loop of chain B (pink) alos stabilize GDP binding. Mg^2+^ is coordinated by four oxygens donated by Ser-157, Glu-242, Asp-193 and an oxygen from the β-phosphate of GDP. mFo-DFc omit map of GDP and Mg^2+^ at 2 σ is shown as a grey mesh.

### MMAA in the M_2_C_2_ complex

Each MMAA monomer in the α_2_ dimer comprises the following signature motifs: switch I, II, and III for signal transduction, a P-loop, base specificity loop and a C-terminal dimerization arm. A structural comparison between free (PDB: 2WWW) and M_2_C_2_- bound (Cα RMSD =3.2 Å) MMAA reveals that one chain undergoes a large inward rotation by ∼60° towards the dimer interface that is driven by an ∼23 Å movement of α-helix7 (residues 296 to 311) in the complex (Figure S1B,C, Movie S2). While the distance between the GDP sites (ribose O’3) in the dimer is unchanged (i.e., 27-28 Å), GDP in the rotated chain moves by 14 Å relative to its position in free MMAA, positioning it closer to the dimer interface and away from solvent (Figure S1C). This open-to-closed motion is more clearly visualized in the surface display of the MMAA structures (Figure 3B,C). The hinge action of the dimerization arm enables the conformational transition from the open to the closed state. A similar albeit larger ∼180° rotation of one subunit of MeaB relative to the other, which buries a previously exposed nucleotide binding site at the dimer interface, was seen in the recent structure of the MeaB-MutAB B_12_ domain complex (Figure S1D) and shortened the distance between the nucleotides from 45 Å to 18 Å in the complex (25). The partially resolved switch I and III loops in free MMAA (23) are resolved in the complex with multiple interactions serving to staple their conformations (Figure S1E, F). Clear electron density is seen for the switch III region in two out of four MMAA chains in the M_2_C_2_ complex, with the switch III loop from chain B interacting with GDP in chain A (Figure S1E). However, the modest side chain electron density of switch III residues, limits accurate modelling of its interaction with GDP. Our model places the side chains of Asp-270 within 4.3 Å to the guanine moiety and Glu-271 within hydrogen bonding distance to the β-phosphate moiety, respectively of GDP (Figure 3D, S1E). Additionally, switch II loops from adjacent subunits interact via the backbone atoms of Val-246 (Figure S1E). The switch III loop was previously captured in three solvent exposed poses in MeaB (15,20). In the MeaB-MutAB B_12_ domain complex, switch III from subunit B makes key contacts with GMPPCP in subunit A via residues that are conserved in MMAA (25) (Figure S2A). Additionally, Lys-276 in MMAA is positioned to serve a similar role as Lys-188 in MeaB, in coordinating to the γ-phosphate in GTP (Figure 3D, S2A).

GDP is anchored via a salt bridge with Asp-292 and hydrogen bonds with Lys-290, Arg-330, Thr-158 and Lys-156, and with backbone interactions with P-loop residues (154-155) as seen previously in the structure of free MMAA bound to GDP (23) (Figure 3D). The GDP site is however, more ordered in the structure of M_2_C_2_ with Mg^2+^ coordinated by three oxygens from the sidechains of Ser-157, Glu-242, Asp-193 and an oxygen from the β-phosphate group of GDP, and two additional water molecules complete the coordination sphere. While the residue corresponding to Asp-180 in MeaB (Asp-92) was predicted to serve as a general base for activating water, its mutation to alanine did not impair the intrinsic GTPase activity (22). In the M_2_C_2_ complex Arg-196 from switch I forms a salt bridge with Glu-271 (Figure 3D). Similarly, in the MeaB-MutAB B_12_ complex, the analogous residue (Arg-108) forms a salt bridge with Asp-182 connecting switch I in chain A to switch III in chain B (Figure S2A). The GDP site in the M_2_C_2_ complex reveals how the product is bound and that conformational changes in switch III would be needed to accommodate the additional phosphate in the substrate, GTP.

### MMUT•MMAA interface

In the M_2_C_2_ complex, the B_12_ domain of MMUT is swapped out by chain A of MMAA, creating a new interface (Figure 4A). An MMUT loop in the substrate domain (spanning residues 227-234), undergoes a large rotation that orients Arg-228 and Tyr-231 for interactions with Asp-344 and Asp-340, respectively on MMAA (Figure S3A). The side chain of Arg-228 is mobile and forms a hydrogen bond with the side chain of the corrin ring in the free MMUT•AdoCbl structure or is rotated away to form a salt bridge with the substrate analog in the MMUT•AdoCbl•malonyl-CoA structure (23). Additional hydrogen bonds that zip this new interface include the backbone carbonyls of Glu-360, Ala-393 and Asp-498 on MMUT interacting with the side chains of Arg-300 and Arg-301 on MMAA, and the side chains of Asp-156 and Gln-476 on MMUT interacting with Ser-336 and Ser-308, respectively (Figures 4B, S4A).

**Figure 4.**
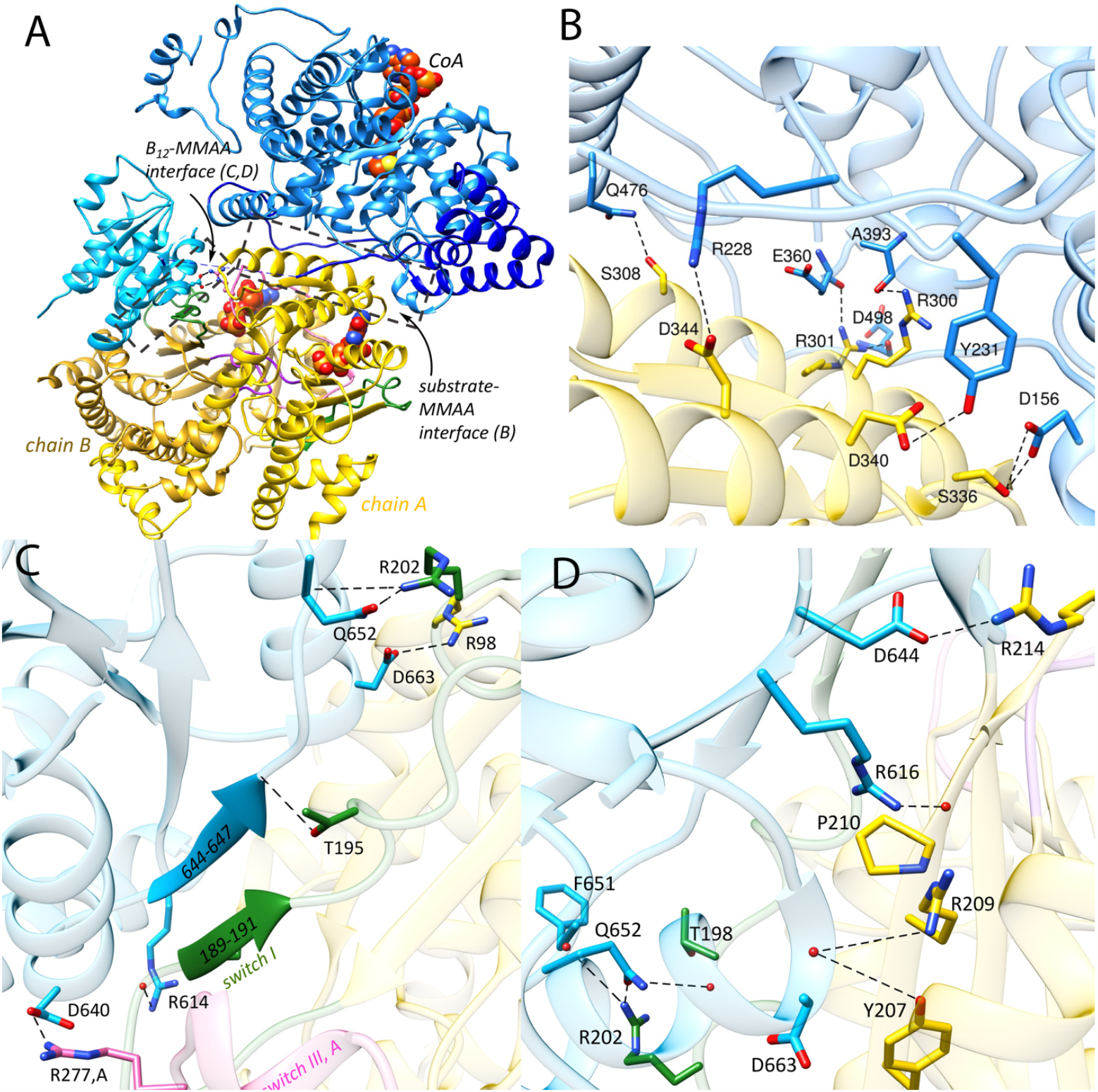
MMUT-MMAA interfaces. **A**. Each MMUT subunit comprising the substrate- (blue) and B_12_ (cyan) domains and the inter-connecting belt (dark blue) in M_2_C_2_ interacts with an MMAA dimer (yellow ribbons). CoA (orange) and GDP (orange) are shown as spheres. Boxes (dashed lines) outline two MMUT-MMAA interfaces. **B**. Close-up of the MMUT substrate domain (blue) and MMAA (chain A, yellow) interface. Black dashed lines represent interactions between residues on MMUT and MMAA (chain A) as described in the text. **C.D**. Close-up of the interface between the B_12_ domain and MMAA (chain B). **C**. Switch I on MMAA (50) is ordered, forming a short β-strand and hydrogen bonds with the terminal β-strand in the B_12_ domain, resulting in an extended β- sheet interaction between the two proteins. Salt bridge and hydrogen bond interactions are shown with dashed lines. **D**. Additional interactions (dashed black line) between MMAA and MMUT residues stabilize the B_12_ domain-MMAA interface as described in the text.

A new interface is also created between the B_12_ domain and MMAA chain B (Figure 4A). Switch I residues (182-198), which are unresolved in free MMAA, are ordered in a short β-sheet (189-191), which hydrogen bonds with the terminal β-strand in the B_12_ domain, creating an extended β-sheet connection between the two proteins (Figure 4C). A similar β-sheet extension from the B_12_- to the G-domain was observed in IcmF and in the MeaB-MutAB B_12_ domain complex (Figure S3B)(18,25). Interestingly, the switch I loop, which links MMAA to the B_12_ domain, also links to switch III in the adjacent MMAA chain (via Arg-196) (Figure 3D.) Although the B_12_ domain interacts primarily with chain B, it also connects via a salt bridge between Asp-640 and Arg-277 to the switch III of chain A (Figure 4C), analogous to the interaction between Asp-609 in the MutAB B_12_ domain and Lys-189 in MeaB (25). The B_12_-MMAA interface is further stabilized by salt bridges between Arg-98 and Asp-663 and Arg-202 and Gln-652 and a hydrogen bond between the backbone atoms of Gly-187 and the side chain of Arg-614 (Figure 4C, S4B). Additionally, the backbone carbonyl of Pro-210 hydrogen bonds with Arg-616 (Figure 4D, S4C). The side chains of Tyr-207 and Arg-209 form hydrogen bonds with the backbone of Asp-663. Arg-214 forms a salt bridge with Asp-644, and Arg-202 and Thr-198 hydrogen bonds with the backbone of Phe-651 and the side chain of Gln-652, respectively.

### MMUT patient mutations disrupt complex formation

*-*The biochemical penalties associated with methylmalonic aciduria causing variants (26-30) that localize at the interfaces between MMAA and the MMUT substrate (R228Q) and B_12_ (R616C and R694W) domains were assessed as a test of the biological relevance of the M_2_C_2_ structure (Figure S5A). The GTPase activating protein (GAP) activity of MMUT serves as a proxy for its complex formation with MMAA. The intrinsic GTPase activity of MMAA (0.06 ± 0.02 min^-1^), which increases 36-fold (2.1 ± 0.2 min^-1^) in the presence of wild-type MMUT, is attenuated 7- (R616C) and 12-fold (R228Q and R694W) in the variants (Table S2). The gating function of MMAA during AdoCbl transfer from MMAB to wild-type MMUT in the presence of GMPPCP was used as an additional test of complex formation (Figure S6A). Complete AdoCbl transfer to R616C MMUT but <10% transfer to R228Q and R694W MMUT was observed, indicating full or partial loss of the GTPase gating function (Figure S6B-E). While AdoCbl-loading from solution was variously impacted by the mutations (Figure S7) the catalytic activity of R228Q was undetectable. The activities of the other variants, R616C (71 ± 4 µmoles min^-1^ mg^-1^) and R694W (73 ± 1 µmoles min^-1^ mg^-1^) were comparable to wild-type MMUT (89 ± 6 µmoles min^-1^ mg^-1^) (Table S2). Cob(II)alamin off-loading from MMUT to MMAB was variously impacted by the mutations; R228Q was unaffected, R694W was partially impaired and R616C was completely impaired (Figure S8). Finally, we tested the impact of simultaneously severing the links between MMAA and the substrate and B_12_ domains of MMUT by engineering the R228Q/R616C double mutant. GAP activation, MMAA-gated AdoCbl loading and cob(II)alamin off-loading functions were all lost (Figures S6,S8). In contrast, AdoCbl binding from solution was unaffected (Figure S7), although catalytic activity was undetectable (Table S2).

### MMAA patient mutations disrupt complex formation

We characterized two MMAA patient mutations, R98G (31) and R209S (32), which reside at the interface with the B_12_ domain in the M_2_C_2_ complex (Figure S5B). While neither mutation affects the intrinsic GTPase activity, GAP activation by MMUT is either enhanced 2-fold (R98G) or lost (R209S) (Table S2). Both mutations impaired nucleotide-gated transfer of AdoCbl from MMAB to MMUT, which was observed even in the presence of GMPPCP (Figure S6) and were unable to power cob(II)alamin off-loading from MMUT for repair (Figures S8). These data are consistent with these disrupting mutations disrupting M_2_C_2_ complex formation.

## Discussion

G-proteins play important roles in metal homeostasis pathways by mechanisms that are generally poorly understood. Structural insights into the mitochondrial G-protein motor MMAA, have been limited by the variable stability and/or oligomerization state of its complex with MMUT, which is influenced by multiple ligands that bind to each protein. In this study, we report the dramatic conformational changes that accompany the M_2_C_2_ nanomotor assembly, exposing the B_12_ domain as a prelude to cofactor translocation (Figure 5).

**Figure 5.**
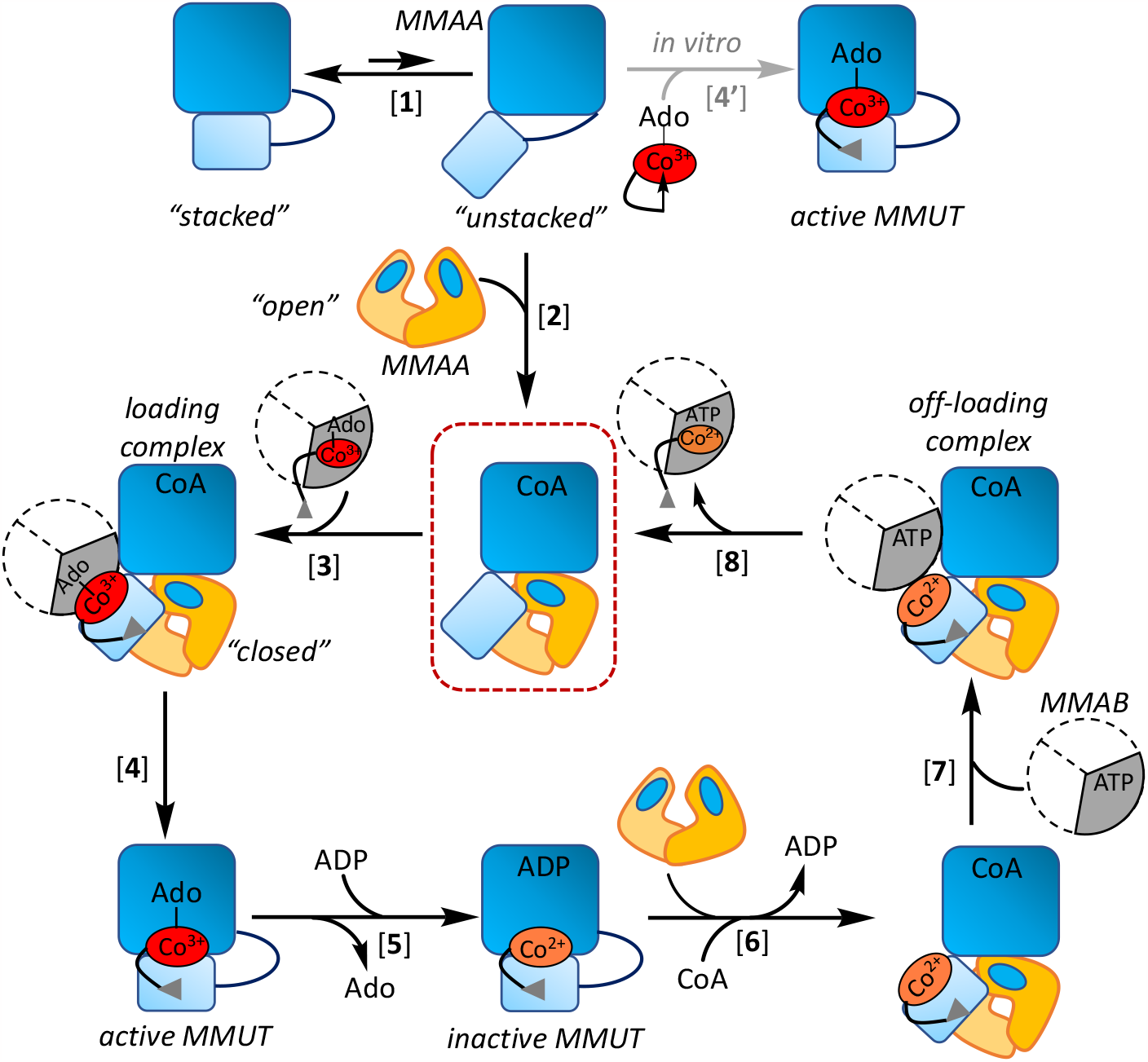
Model of nanomotor assemblies involved in B_12_ loading and repair. Free MMUT (light and dark blue are the B_12_ and substrate domains) exists in the “stacked” and “unstacked” conformations with the equilibrium favoring the former [1]. MMAA (yellow) stabilizes the unstacked conformation by wedging between the substrate and B_12_ domains and itself transitions from an “open” to “closed” state in M_2_C_2_, crystallized in this study (red box) [2]. MMAB transfers AdoCbl to M_2_C_2_ [3], leading to catalytically active holo-MMUT [4]. *In vitro*, free AdoCbl can bind to MMUT [4’]. Inactivation of MMUT due to loss of Ado (deoxyadenosine) followed by ADP binding, stabilizes cob(II)alamin against hyperoxidation [5] and leads to recruitment of the repair complex [6] and [7]. Following cob(II)alamin off-loading and conversion to AdoCbl by MMAB, the active cofactor is reloaded onto MMUT [8].

In all previous structures of MMUT and its bacterial orthologs, the B_12_ and substrate domains are stacked on each other, forming a solvent-protected active site (23,33,34). The apo- and inactive forms of MMUT (with cob(II)alamin bound) exhibit a higher affinity for MMAA and the M_2_C_2_ structure explains how its assembly harnesses the GTPase activity to make the B_12_ domain accessible for cofactor transfer. Apo-MMUT can be reconstituted with free AdoCbl *in vitro*, suggesting that the unstacked conformation of MMUT is available even in the absence of MMAA (Figure 5). However, this path is unlikely to be physiologically relevant since the cellular concentration of free AdoCbl is negligible (35). MMAA wedges physically between the two MMUT domains and stabilizes a heretofore unseen conformation in which the B_12_-domain is swung out by 180° degrees relative to the substrate domain, exposing the highly conserved His-627 cobalamin ligand to solvent. Mutation of the corresponding histidine ligand in MutAB leads to AdoCbl loading failure, revealing a critical role for the thermodynamically favored increase in coordination number (from 5- to 6-coordinate) for cofactor translocation from MMAB to MMUT (7) Crystallographic snapshots of the *Mycobacterium tuberculosis* MMAB ortholog captured multiple poses that provide clues into how AdoCbl, but not cob(II)alamin, is selectively translocated to the mutase (36).

Once loaded with AdoCbl, the affinity of holo-MMUT for MMAA is significantly weakened (8), leading us to propose that the catalytically active form of the mutase functions as a stand-alone dimer (Figure 5). Inactivation of MMUT with resultant cob(II)alamin accumulation, necessitates cofactor repair with concomitant protection against hyperoxidation to the aquo-cob(III)alamin state, which represents a dead-end complex. We have recently discovered bivalent mimicry by ADP, an abundant metabolite, induces a conformational changes that protects cob(II)alamin from over-oxidation (37). Cob(II)alamin off-loading to MMAB from this state is promoted by CoA (or M-CoA) binding to MMUT and by MMAA-dependent GTP hydrolysis (Figure 5) (37). It is presently unclear however, how inactive MMUT (with cob(II)alamin and CoA) is distinguished from the active enzyme (with AdoCbl and M-CoA) to recruit the repair system (MMAB and MMAA).

Free MMAA exhibits an “open” conformation in which the nucleotide binding site is only partially ordered, explaining its low intrinsic GTPase activity (23). In the M_2_C_2_ complex, MMAA is in a “closed” conformation, which brings its active site into register, explaining the GAP function of MMUT (Figure 3C,D). Each switch III domain in the MMAA dimer, previously shown to be important for its GTPase activity (8,20), completes the nucleotide binding site in the opposite monomer in the M_2_C_2_ complex. A similar switch III crossover strategy for building the active site was seen in the recent MeaB-MutAB B_12_ domain structure (25). Comparable switch III interactions are however, absent in IcmF where the two G-domains are at opposite ends of the protein (Figure S2B) (18). G-domain dimerization would require the linear association of two IcmF dimers. Formation of an active nucleotide binding site at the dimer interface was also observed in HypB, a G-protein involved in nickel trafficking, where residues from one chain contribute to nucleotide binding in the adjacent chain (38). Interestingly, the interprotein β-sheet extension between switch I and the B_12_ domain is a structural innovation that recurs in IcmF(18), MeaB-MutAB B_12_ domain (25), and in the human M_2_C_2_ complex (Figure S3B). This motif confers rigidity to the interface and provides a direct conduit for signal transmission from the nucleotide to the B_12_ site.

The unstacked conformation provides the first structural insights into how MMAB might access the B_12_-binding site in the M_2_C_2_ complex and the role of MMAA in facilitating this process (Figure 5). We speculate that the GDP and CoA bound M_2_C_2_ complex captured in this study, could also serve a regulatory role for sequestering MMUT and prioritizing B_12_ for the cytoplasmic branch (39). In liver, ∼25% of MMUT is B_12_ loaded, whereas cytoplasmic methionine synthase exists predominantly in the holo-form (40,41). MMAA is predicted to be predominantly GDP-loaded based on the intracellular concentrations of GDP (160 μM) and GTP (∼470 μΜ) relative to their respective *K*_D_ values for MMAA (1.1 μΜ (GDP) and 740 μΜ (GTP)) (8) Apo-MMUT is likely to predominantly CoA bound based on the *K*_D_ (113 µM) versus the mitochondrial concentration of CoA (2-5 mM) (37) Thus, the M_2_C_2_ complex captured here could represent a holding structure that would be primed for loading upon exchange of GDP with GTP and availability of AdoCbl-bound MMAB.

The dramatic stacked to unstacked conformational change could be a common strategy used by B_12_-dependent enzymes, particularly those that bind the cofactor in the base-off conformation. Ligand-triggered conformational changes have been reported in bacterial methionine synthase, although the mechanism of cofactor loading awaits elucidation (42). In *Aquincola tertiaricarbonis*, a MeaB like gene is in the same operon as the large subunit of cobalamin-dependent hydroxyisobutyryl-CoA mutase (43). Although the substrate (large) and B_12_ (small) subunits are on separate polypeptides, the mechanism of B_12_ loading onto this mutase could represent a variation on the same theme as in MMUT. On the other hand, AdoCbl-dependent diol dehydratase (44), glycerol dehydratase (45), ethanolamine ammonia lyase (46) and glutamate mutase (47) use an ATP dependent reactivase to repair the inactive cofactor. Exchange of ADP with ATP releases the activase and frees up the apo enzyme for AdoCbl loading (48). A docking model of diol dehydratase with its reactivase suggests that the substrate domain must tilt relative to the B_12_ domain to avoid steric clashes in the complex (49).

In summary, the structure of the M_2_C_2_ motor provides insights into a common intermediate in the bidirectional movement of the B_12_ cofactor between human MMUT and MMAB (Figure 5). The dramatic change in solvent access of the B_12_-domain suggests that it serves as a platform for recruiting MMAB. MMAA also undergoes large conformational changes that complete its active site architecture and explains the molecular basis of the GAP activity of MMUT. Patient mutations at the MMAA-MMUT interfaces impact complex formation and highlight the importance of protein dynamics in translocating a large cofactor.

## Author Contributions

R.M., and M.Y. established crystallization conditions and R.M. elucidated the crystal structure. M.R, H.G. and N.H. designed and performed the biochemical analyses. All authors helped conceive the experiments and analyze data. RM, MR and RB drafted the manuscript, and all co-authors edited it.

## Data and materials availability

All data are available in the manuscript or supplementary materials. The structure factors and coordinates for human MMUT•MMAA•CoA•GDP (PDB code: 8GJU) have been deposited in the Protein Data Bank.

## Acknowledgement

RM thanks Dr. Dali Liu (Loyola University Chicago) for helpful discussions on structure refinement.

## Supplementary Materials

Materials and Methods

Tables S1 and S2

Figures S1 to S9

Movie S1 and S2

