## supporting information for "Architecture of the human G-protein-methylmalonyl-CoA mutase nanoassembly for B_12_ delivery and repair"

### Methods

#### *Materials*

Adenosine 5'-triphosphate disodium salt hydrate (ATP) (Cat. # A2383), Coenzyme A disodium salt (CoA) (Cat. # C3144), guanosine 5'-triphosphate sodium salt hydrate (GTP) (Cat. # G8877), guanosine 5'-diphosphate sodium salt (Cat. # G7127), β,γ-methyleneguanosine 5`-triphosphate sodium salt (GMPPCP) (Cat. # M3509), and 5'deoxyadenosine cobalamin (AdoCbl) (Cat. # C0884) are from Sigma-Aldrich. Isopropyl $\beta$-D-1-thiogalactopyranoside (IPTG) (Cat. # I2481C) and Tris (2-carboxyethyl) phosphine (TCEP) (Cat. # TCEP) are from Gold Biotechnology. Ni(II)-NTA resin (Cat. # 30210) was from Qiagen. Primers were purchased from Integrated DNA Technologies. Cob(II)alamin was prepared by photolysis of AdoCbl as described previously (1).

#### *Expression and purification of wild-type and mutant MMUT*

The following forward primer sequences were used for generating the, R228Q, R616C, R694W and R228Q/R616C MMUT mutants using the wild-type MMUT expression clone in a pET-28b vector with a C-terminal TEV cleavable His-tag. The reverse primer had a complementary sequence.

R228Q:5'GATATTCTGAAAGAATTTATGGTTCAAAACACCTACATCTTCCCGCCGGAACCG-3'

R616C: 5`CGTCGCCCGTGCCTGCTGGTTGCCAA3`

R694W: 5`GAACTGAACAGCCTGGGCTGGCCGGATATTCTGGTCATGTGT3'

The mutations were verified by Sanger nucleotide sequence analysis (Eurofins). Recombinant wild-type and R616C and R694W MMUT were expressed as reported earlier (37). The R228Q and R228Q/R616C variants were expressed at 12 ˚C following induction. Wild-type and mutant

MMUT were purified as described previously (2) and were obtained in yields of: 15 mg (wild-type), 7 mg (R228Q), 9 mg (R228Q/R616C), 10 mg (R616C), and 4 mg (R694W) per liter of culture.

***Expression and purification of MMAB and MMAA***

Wild-type and mutant MMAA, and wild-type MMAB were purified as reported (2) The MMAA mutants R98G and R209S were generated by the Quickchange protocol using the following forward primers. The reverse primer had a complementary sequence.

R98G: 5'-CCGGTCTGATTCAAGGTCAAGGTGCGTGCCTG-3'

R209S: 5'-GTGACATGAACGCGTACATTAGTCCGAGCCCGACCCG-3'

***AdoCbl loading to MMUT from MMAB***

In a quartz cuvette, 15 μM AdoCbl and 15 μM MMAB in Buffer A (50 mM HEPES pH 7.5, 150 mM KCl, 2 mM $MgCl_2$, 2 mM TCEP and 5% glycerol) were incubated for 5 min at 25 ºC to prepare MMAB•AdoCbl. AdoCbl transfer was initiated by addition of a premixed solution of 15 μM MMUT 30 μM MMAA, 1 mM GMPPCP to the reaction mixture at 25ºC. The spectrum of AdoCbl was monitored between 300 and 700 nm and recorded every 60 sec for 30 min at 25 ºC. Transfer of AdoCbl from MMAB to MMUT is signaled by an increase in absorbance at 525 nm.

***Cob(II)alamin off-loading from MMUT to MMAB***

Cob(II)alamin off-loading from MMUT was monitored under anaerobic conditions (Omni-Lab anaerobic chamber, containing < 0.5 ppm $O_2$) in a quartz cuvette containing Buffer A, 15 μM cob(II)alamin and 10 μM MMUT, incubated for 10 min at 20 ºC. The "repair mixture" was prepared by mixing 5 mM ATP, 15 μM of MMAB, 30 μM MMAA and 1 mM GTP. Cob(II)alamin transfer was initiated by adding the repair mixture to the cuvette at 20ºC. The spectrum was recorded between 300 and 750 nm every 15 sec for 15 min. Transfer of cob(II)alamin from MMUT to MMAB is signaled by an increase in absorbance at 464 nm, which corresponds to 4-coordinate cob(II)alamin bound to MMAB.

***MMUT GAP activation of MMAA***

To measure the GAP activation by MMUT variants, samples were prepared by mixing 2.5 μM MMAA and 2.5 μM wild-type or mutant MMUT in 50 mM HEPES pH 7.5, 150 mM KCl, 2 mM $MgCl_2$, 2 mM DTT and 5% glycerol in a 1.5 mL Eppendorf tube, and pre-incubated at 30 °C for 10 min. The intrinsic GTPase activity of wild-type, R98G and R209S MMAA GTPase was measured using 12.5 μM of each protein. The reaction was initiated by adding 3 mM GTP to the Eppendorf tubes at 30 °C. Aliquots of 50 μL were removed after 10 min and 20 min and the reaction was immediately terminated by adding 2.5 μL of 2M trichloroacetic acid and placed on ice. The precipitant was removed by centrifugation at 15,870 x *g* for 10 min at 4 °C and the GDP concentration in the supernatant was analyzed by HPLC as described previously (1). The concentration of GDP present in the control sample (lacking proteins) was subtracted from each value. A calibration curve for GDP (0-500 μM) was obtained by treating standard samples as mentioned above.

***Catalytic activity of MMUT***

MMUT activity was assessed in the thiokinase-coupled spectrophotometric assay as described previously (2). A stock solution of holo MMUT was prepared by mixing 10 μM MMUT with 20 μM AdoCbl in 100 mM potassium phosphate, 3 mM $MgCl_2$ pH 7.5 buffer followed by incubation at 30 °C for 15 min. In a quartz cuvette, 5 μM AdoCbl, 3 mM GDP, 5 μM thiokinase, 0.5 mM M-CoA, and 70 μM 5,5´-dithiobis(2-nitrobenzoic acid) was incubated at 30 °C in 100 mM potassium phosphate, 3 mM $MgCl_2$ pH 7.5 buffer in a final volume of 200 µL and the background thioesterase activity was recorded at 412 nm for 5 min. The reaction was initiated by addition of 2 µL of holo-MMUT stock (2 nM final concentration) to the cuvette. The specific activity was calculated assuming that the amount of CoA detected was directly proportional to the concentration of succinyl-CoA formed by MMUT. An extinction coefficient of 14,150 $cm^{-1}$ $M^{-1}$ was used for the $TNB^{-}$ anion (3).

#### *Crystallography*

The $M_2C_2$ complex with GDP and CoA was prepared by mixing 100 µM MMUT, 200 µM MMAA, 2 mM GDP and 2 mM CoA in 50 mM HEPES, pH 7.5, 75 mM KCl, 2 mM $MgCl_2$, 2 mM TCEP. The mixture was purified on a Sephacryl S300 16/600 column (Cytiva) pre-equilibrated with the same buffer (Figure S9). Fractions (2.5 mL) were collected in tubes containing 2.5 µL 100 mM GDP and 2.5 µL 100 mM CoA (final concentration 100 µM each). The $M_2C_2$ complex-containing fractions were pooled and concentrated to ~30 mg/mL. GDP and CoA were added to a final concentration of 500 µM each. The complex was flash frozen in liquid nitrogen and stored at -80 ˚C. Crystals of $M_2C_2$ appeared within a day using the hanging drop vapor diffusion method. Crystals with the best diffraction were obtained in a 1 µL drop containing 1:1 protein (15 mg/mL): well solution (Morpheus 1 F11 (Molecular Dimensions): 120 mM monosaccharides mix, 100 mM buffer system 3, pH 8.5, 30% precipitant mix 3) (4). Crystals were harvested after a week and flash cooled in liquid nitrogen for data collection. No additional cryoprotectant was used.

#### *Data collection, processing, and refinement*

Diffraction data were collected at GMCA (23-ID-B) at Argonne National Laboratory. X-ray diffraction data were processed with autoPROC, using the default pipeline which includes XDS,

Truncate, Aimless, and STARANISO (Tickle et al. STARANISO, 2018, Global Phasing Ltd, UK) (5-7). Analysis by STARANISO showed that diffraction data were anisotropic, with diffraction limits along the reciprocal directions of 2.79 Å along 0.967*a** – 0.253*c**, 3.62 Å along *b**, and 3.15 Å along -0.05 *a**+0.999*c**. Automated resolution cutoff with local $I/\sigma(I)$ > 1.20 of anisotropic corrected data by STARANISO resulted in a 2.79 Å resolution data set. Data processing statistics are shown in Table S1.

The structure was solved by molecular replacement with Phaser (8), using previously solved structures of MMUT (PDB code 2XIQ) and MMAA (PDB code 2WWW). The $M_2C_2$ crystals belonged to the space group P 21 with 4 chains of MMUT and MMAA per asymmetric unit. Iterative rounds of model building, and refinement were performed with COOT(9) and refined in Phenix (10) or Refmac (6). Ligand restraints were generated in *eLBOW* (11). The geometric quality of the model was assessed in *MolProbity* (12). Models were analyzed in Pymol (Schrödinger, LLC) and the structure figures were generated using UCSF Chimera (13). Model refinement statistics are shown in Table S1.

**Table S1.** Crystallographic data for the human $M_2C_2$ complex

| | MMUT•MMAA•GDP•CoA |
|---|---|
| Beamline | APS, GMCA-B |
| Wavelength (Å) | 1.033 |
| Temperature (K) | 100 |
| Space group | P 1 21 1 |
| Cell dimension | |
| α, β, γ (°) | 90, 105.5, 90 |
| a, b, c (Å) | 95.8, 221.8 121.8 |
| Resolution (Å) | 117.3-2.79 (3.07-2.79) |
| $R_{merge}$ (%) | 13.7 (77) |
| $R_{meas}$ (%) | 16.6(92) |
| $R_{pim}$ (%) | 9.2 (50) |
| <I/σ> | 5.2 (1.6) |
| CC (½) | 1.0 (0.73) |
| Completeness (%) (spherical) | 61.9(9.1) |
| Completeness (%) (ellipsoidal) | 89.3(63.9) |
| Multiplicity | 3.1 (3.1) |
| No. Reflections | 234358(8485) |
| No. Unique Reflections | 7483 (2737) |
| Overall B (Å$^2$) (Wilson plot) | 71.9 |
| Resolution Range | 80.6-2.79 (2.83-2.79) |
| Number of reflections (work/test) | 74782/3731 |
| $R_{work}$ / $R_{free}$ (%) | 0.23/0.27 |
| No. of atoms | |
| protein | 30,235 |
| water | 14 |
| Ligand: CoA | 192 |
| GDP | 112 |
| B-factors(Å$^2$) | |
| protein | 63.82 |
| water | 48.57 |
| Ligand: CoA | 51.9 |
| GDP | 53.2 |
| Rmsd deviations | |
| Bond lengths (Å) | 0.004 |
| Bond angles (°) | 0.633 |
| Ramachandran plot (%) Favored, allowed, outliers | 96.04, 3.96, 0 |
| MolProbity score | 2.05 |
| PDB code | 8GJU |

**Table S2:** Summary of the biochemical penalties associated with MMUT and MMAA variants[1]

| | GTPase activity (min$^{-1}$) | Intrinsic GTPase activity (min$^{-1}$) | Fold change | Activity (μmoles min$^{-1}$ mg$^{-1}$) | AdoCbl transferred | Cob(II)alamin transferred |
|---|---|---|---|---|---|---|
| wild type MMUT | 2.1 ± 0.2 (n = 3) | N/A | 36 | 89 ± 6 (n = 3) | blocked | complete |
| R228Q | 0.7 ±0.1 (n = 4) | N/A | 12 | ND (n = 3) | partial | complete |
| R616C | 0.4 ± 0.1 (n = 3) | N/A | 7 | 71 ± 5 (n = 3) | complete | blocked |
| R694W | 0.7 ± 0.3 (n = 3) | N/A | 12 | 73 ± 1 (n = 3) | blocked | partial |
| R228Q/R616C | ND (n = 3) | N/A | ND | ND (n = 3) | complete | partial |
| R98G MMAA | 0.08 ± 0.05 (n = 4) | 0.05 ± 0.03 (n = 6) | 2 | N/A | complete | blocked |
| R209S MMAA | 0.03 ± 0.01 (n = 4) | 0.03 ± 0.02 (n=6) | 1 | N/A | complete | blocked |

[1]The GTPase activity of MMAA in the presence of stoichiometric MMUT was measured. The fold change was calculated by dividing the GTPase activity in the presence of MMUT by the intrinsic GTPase activity of wild-type MMAA (0.06 ± 0.02 (n = 8)) or the MMAA variants in the absence of MMUT. n = number of repeats, ND = not detected, N/A = not applicable.

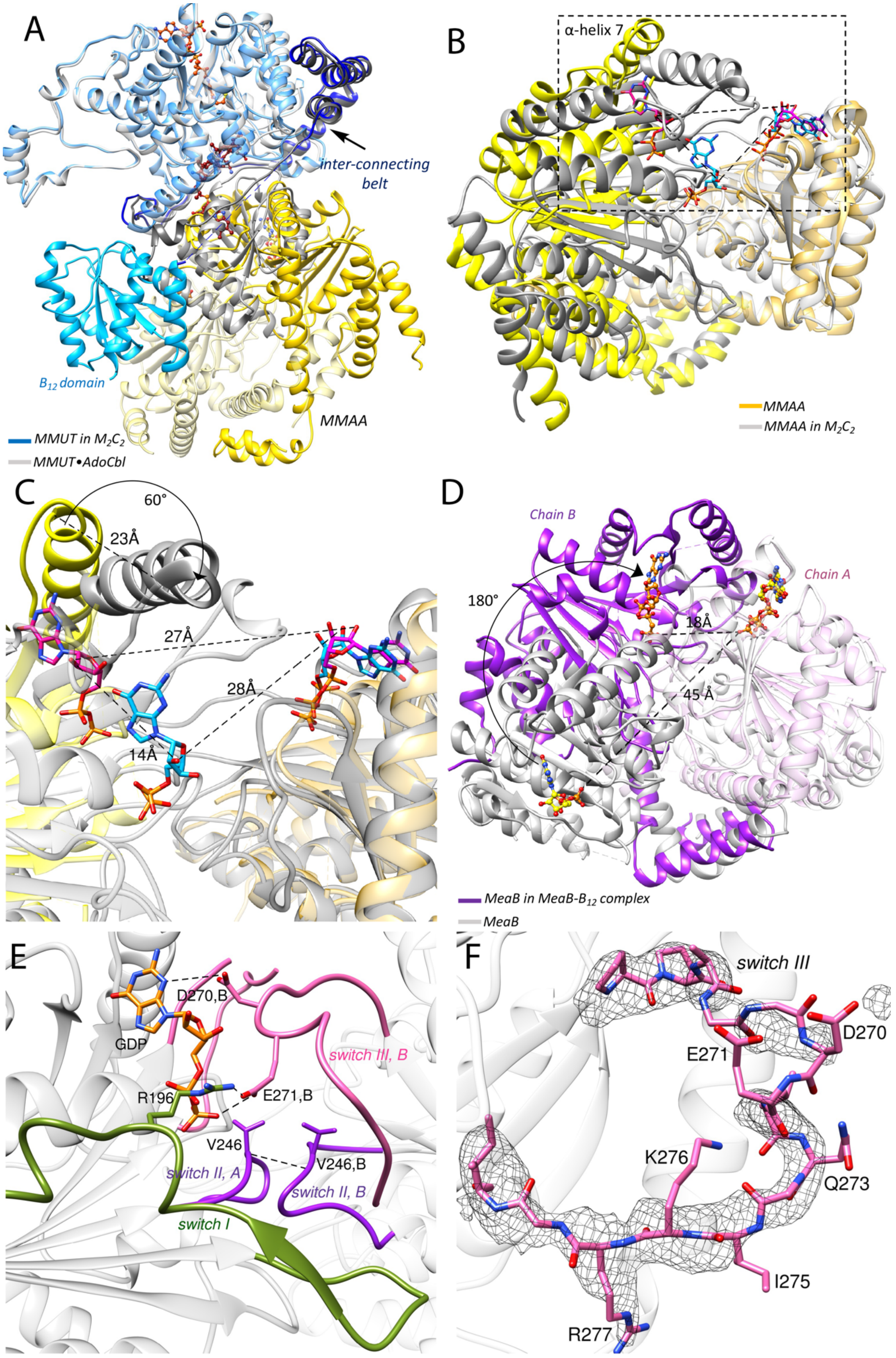
A
inter-connecting belt
$B_{12}$ domain
MMAA
MMUT in $M_2C_2$
MMUT•AdoCbl
B
α-helix 7
MMAA
MMAA in $M_2C_2$
C
60°
23Å
27Å
28Å
14Å
D
Chain B
Chain A
180°
18Å
45 Å
MeaB in MeaB-$B_{12}$ complex
MeaB
E
D270,B
GDP
switch III, B
R196
E271,B
V246
V246,B
switch II, A
switch II, B
switch I
F
switch III
D270
E271
K276
Q273
I275
R277

**Figure S1. A.** Overlay of MMUT in $M_2C_2$ (blue ribbon) and MMUT·AdoCbl·malonyl-CoA (PDB: 2XIQ) shows the $B_{12}$ domain becomes solvent exposed in $M_2C_2$ (cyan). MMAA (yellow ribbons) wedges between the $B_{12}$ and substrate domains, forming new protein-protein interfaces. AdoCbl (red) and CoA (orange) are shown in stick display. **B.** Overlay of MMAA in the $M_2C_2$ complex (grey) with free MMAA (PDB: 2WWW) (yellow). **C.** A close-up of the boxed area in B shows that one subunit rotates by ~60° towards the dimer interface. GDP is shown in blue stick display in $M_2C_2$ and in magenta in free MMAA. **D**. Overlay of free MeaB (PDB:2QM7, grey ribbon) and MeaB in complex with the MutAB $B_{12}$ domain (PDB:8DPB). The purple ribbon shows that chain B undergoes a 180° rotation upon complex formation. GDP in MeaB is shown as yellow sticks and GMPPCP in the MeaB-$B_{12}$ domain is shown in orange. **E.** A close-up of the MMAA dimer interface shows that switch I (green), switch II (purple) and switch III from chain B (pink) are engaged in GDP (orange sticks) binding in the $M_2C_2$ complex. **F**. A 2mFo-DFc composite omit map at 1.5 σ of the switch III loop is shown in grey mesh. The backbone and side chains atoms of residues in the switch III loop are shown in pink stick display.

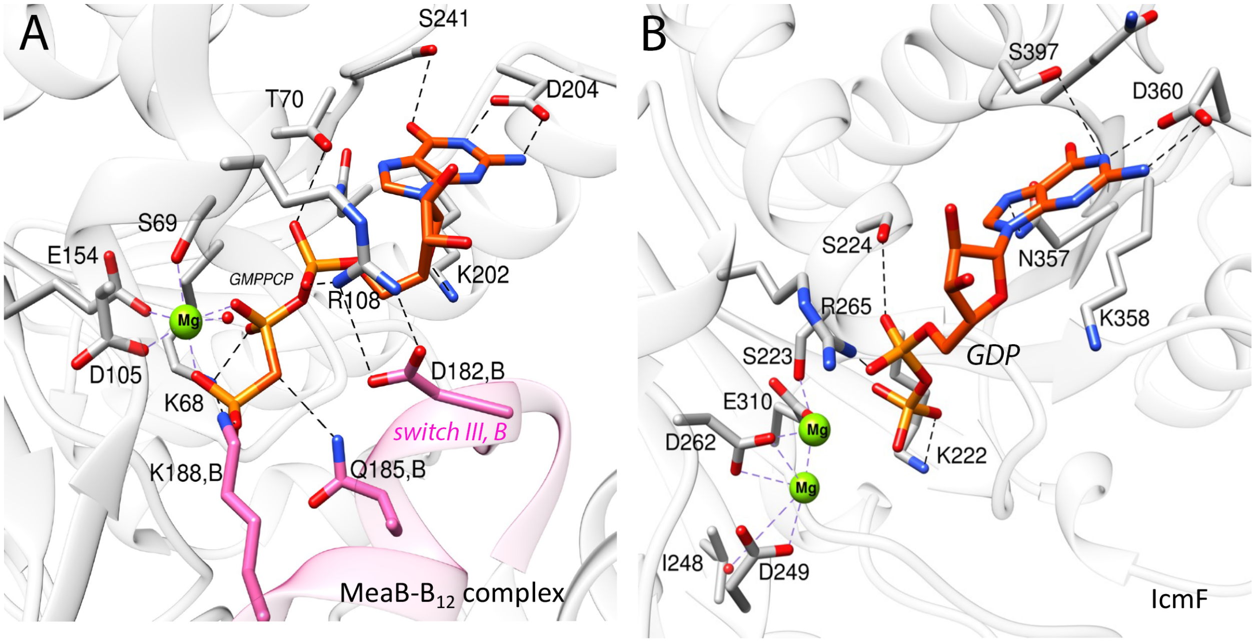


**Figure S2**. **A**. Nucleotide binding site in the MeaB-MutAB $B_{12}$ domain complex. $Mg^{2+}$ (green sphere) is coordinated by oxygens from the α- and β-phosphate groups in GMPPCP (orange sticks), the oxygens from the side chains of Ser-69, Glu-154, Asp-105, and a water molecule. Switch III residues (pink) Gln-185 and Lys-188 from chain B form hydrogen bonds with GMPPCP, and Asp-182 forms a salt bridge with Arg-108. The corresponding residues in human switch III are Asp-270, Gln-273 and Lys-276. **B** The nucleotide binding site in IcmF. GDP is shown in orange sticks. Two $Mg^{2+}$ ions (green spheres) are coordinated by Ser-223, Glu-310, Asp-262, and Asp-249.

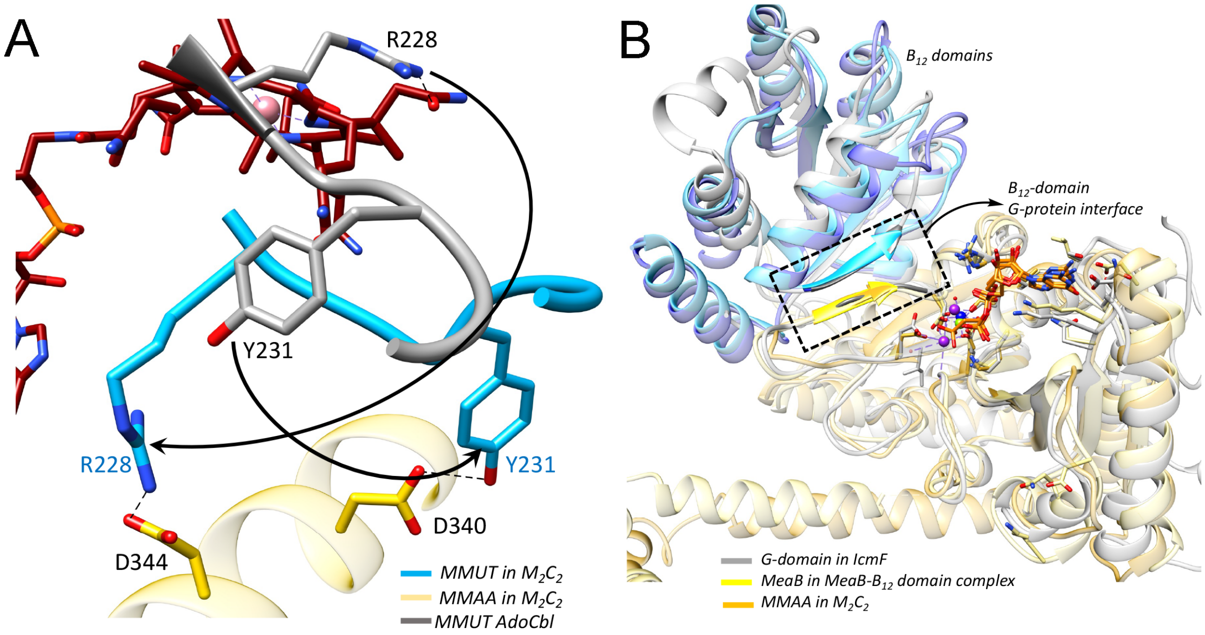


**Figure S3. A**. Overlay of MMUT•AdoCbl (grey, PDB:2XIQ) and MMUT in the $M_2C_2$ complex (blue) showing reorientation of a loop (227-234), which positions Arg-228 and Tyr-231 for interactions with Asp-344 and Asp-340 in MMAA. Cobalamin is shown as red sticks. **B**. Overlay of the $B_{12}$-(blue) and G- (yellow) domains in $M_2C_2$ with the corresponding domain in IcmF (grey, PDB: 4XC8) and the MeaB-MutAB $B_{12}$ domain complex (PDB:8DPB) reveals conservation in their juxtaposition. The boxed region highlights the β-sheet extension between switch I and the terminal β-sheet in the $B_{12}$-domain. GDP or GMPPCP is shown as orange sticks. Mg is shown as green, purple or dark blue spheres in MMAA, IcmF and MeaB, respectively.

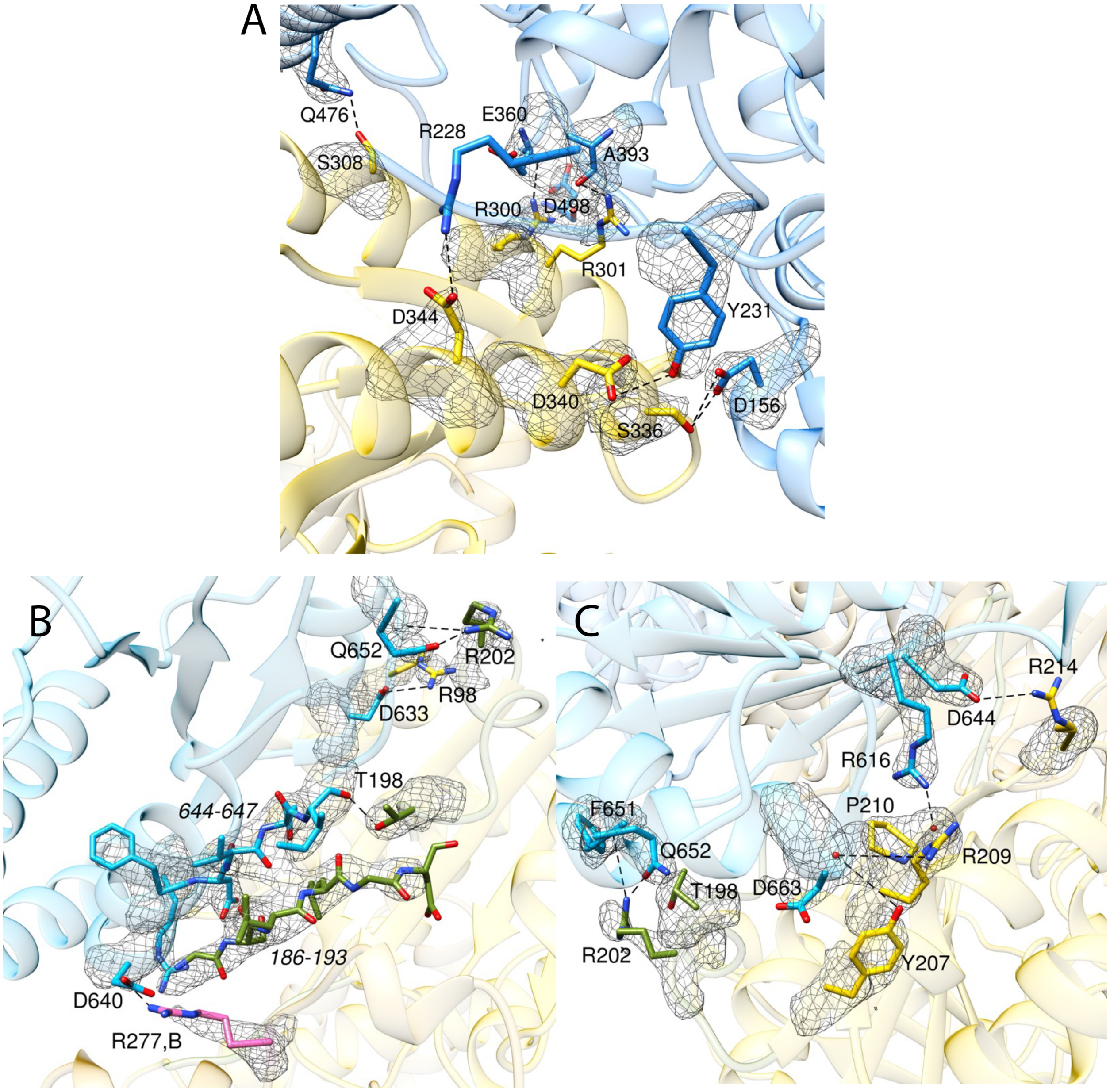


**Figure S4. A** Residues interacting at the substrate domain (blue)-MMAA (yellow) interface are shown as sticks. 2mFo-DFc composite omit map of these residues is shown as a grey mesh at 1.5 σ. **B.C.** Residues interacting at the $B_{12}$ domain (blue)-MMAA (yellow) interface and a 2mFo-DFc composite omit map of these residues is shown as a grey mesh at 1.5 σ. Residues on the $B_{12}$-domain and MMAA are shown as blue and yellow sticks, respectively, Switch I residues are shown as green sticks and Arg-277 from switch III is in pink.

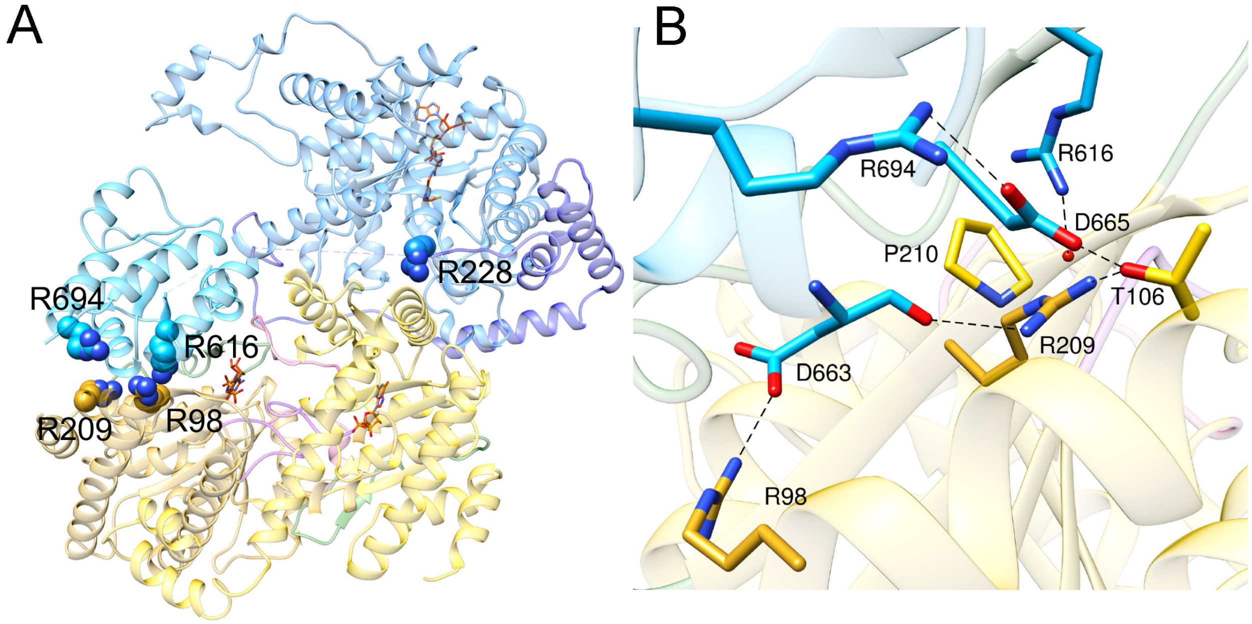


**Figure S5.** Patient mutations at the MMUT•MMAA interfaces. **A.** Arg-228 resides at the substrate domain-MMAA interface while Arg-616 and Arg-694 in the $B_{12}$-domain are involved in a different interface with the second subunit of MMAA. Arg-98 and Arg-209 from MMAA also reside at the $B_{12}$-domain-MMAA interface. **B.** Close-up of the $B_{12}$ domain-MMAA interface showing interactions between Arg-616 and the backbone of Pro-210. The sidechain of Asp-665 bridges Arg-694 to MMAA via a salt bridge with Arg-694 and a hydrogen bond to Thr-106. Arg-209 and Arg-98 form hydrogen bonds with the backbone and sidechain of Asp-663.

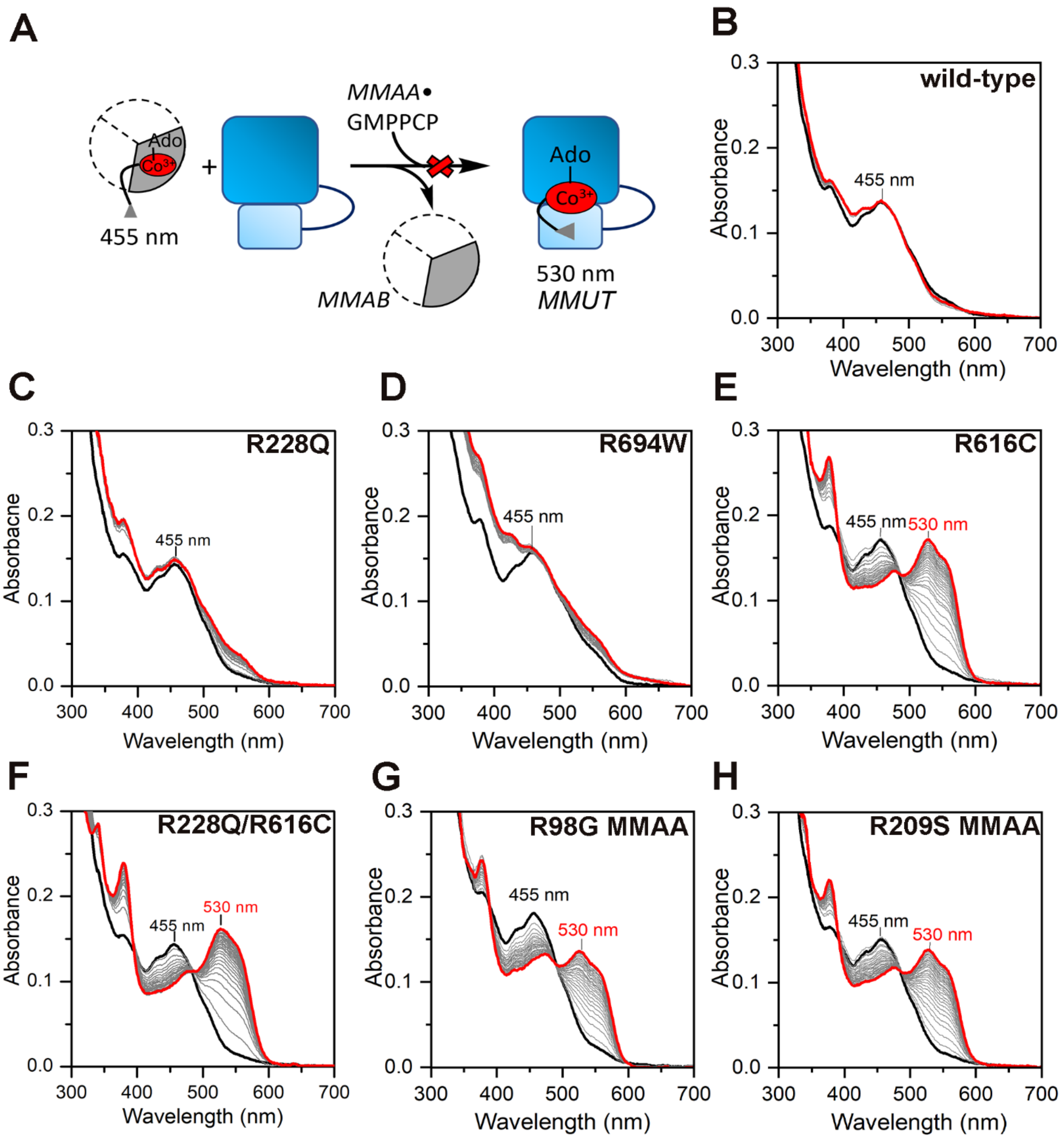


**Figure S6. AdoCbl transfer from MMAB to MMUT in the presence of MMAA•GMPPCP**. (A) Scheme showing the gated transfer of AdoCbl from MMAB to MMUT, which does not occur when the non-hydrolysable GTP analogue GMPPCP is bound to MMAA. When mutations disrupt $M_2C_2$ complex formation, AdoCbl transfer from MMAB to MMUT is accompanied by a spectroscopic shift from 455 nm (MMAB•AdoCbl) to 530 nm (MMUT•AdoCbl). (B -H) Transfer of AdoCbl (15 µM) bound to MMAB (15 µM) in Buffer A following addition of wild-type or mutant MMUT (15 µM), wild-type or mutant MMAA (30 µM) and GMPPCP (1 mM). UV/visible spectra were recorded every minute for 30 min. The initial and final spectra are shown in black and red respectively. (**B**) wild-type MMUT, (**C**) R228Q MMUT, (**D**) R694W MMUT (**E**) R616C MMUT (**F**) R228Q/R616C MMUT, (**G**) R98G MMAA, and (**F**) R209S MMAA. The spectra are representative of three independent experiments.

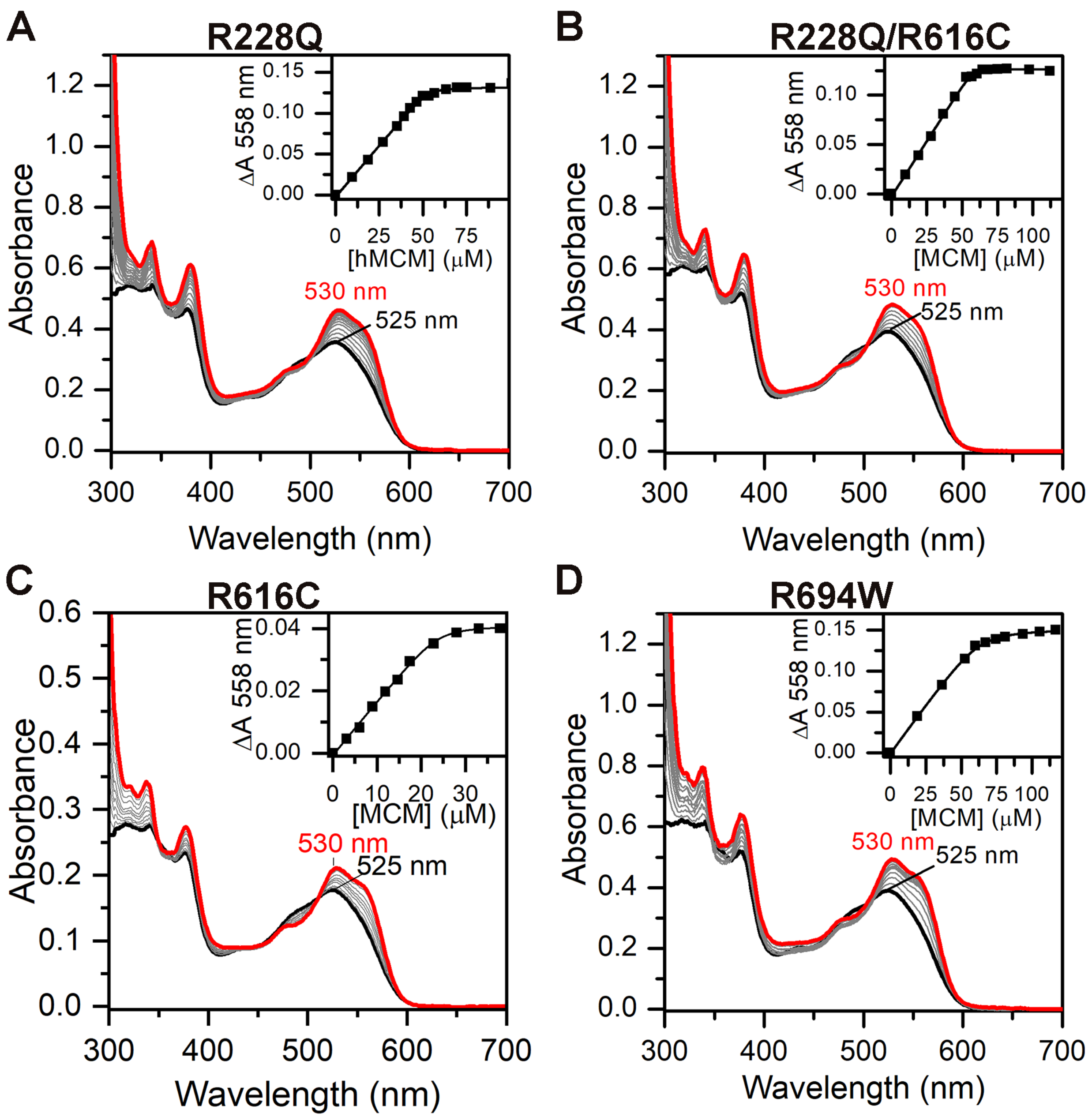


**Figure S7. Determination of dissociation constants for binding of free AdoCbl to MMUT**. Varying concentrations of the MMUT variants were titrated into a solution of AdoCbl (50 μM) in Buffer A at 25 ˚C. (**A**) R228Q MMUT, (**B**) R228Q/R616C MMUT, (**C**) R616C MMUT, and (**D**) R694W MMUT. *Insets*. Dependence of $A_{558nm}$ on MCM concentration. The data are representative of three independent experiments. Fitting of the binding isotherms in the insets to a single site binding model yielded the following $K_{D(AdoCbl)}$ values: wild type, R228W and R228W/R6161C =<0.2 μM, R6161C= 0.5 ± 0.2 μM, and R694W (1.1 ± 0.1 μM).

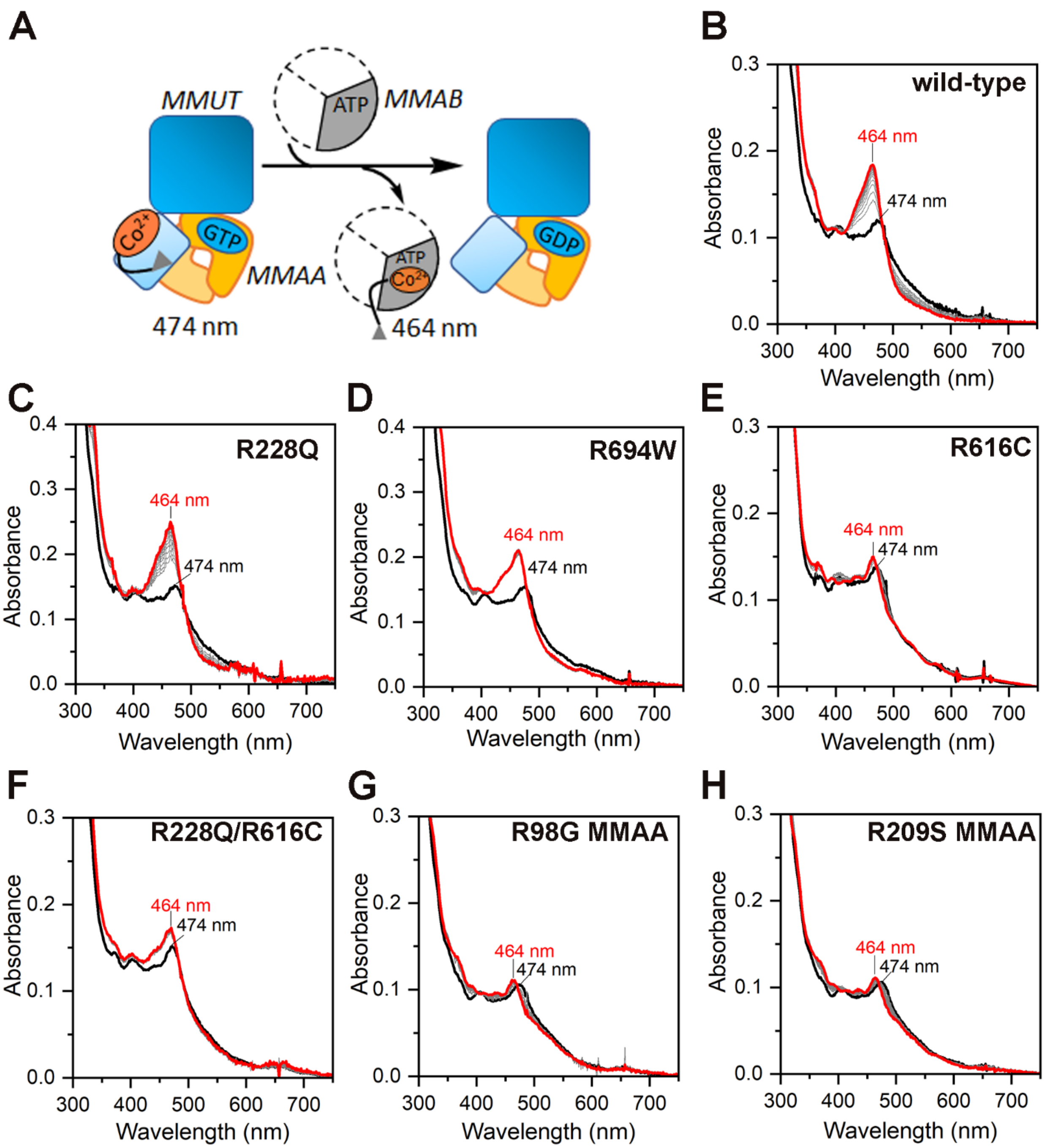


**Figure S8. Cob(II)alamin transfer from MMUT to MMAB.** (**A**) Scheme showing the transfer of cob(II)alamin from MMUT to MMAB in the presence of MMAA•GTP. The blue shift from 474 nm to 464 nm with a concomitant increase in intensity is indicative of successful transfer. (**B-H**) Transfer of cob(II)alamin (15 μM) bound to wild-type or mutant MMUT (10 μM) in Buffer A following addition of MMAB (15 μM), wild-type or mutant MMAA (30 μM), 1 mM GTP and 5 mM ATP. UV/visible spectra were recorded every minute for 15 min; the initial and final spectra are shown in black and red, respectively. (**B**) Wild-type MMUT, (**C**) R228Q MMUT, (**D**) R694W MMUT, (**E**) R616C MMUT, (**F**) R228Q/R616C MMUT, (**G**) R98G MMAA, and (**H**) R209S MMAA. The spectra are representative of three independent experiments.

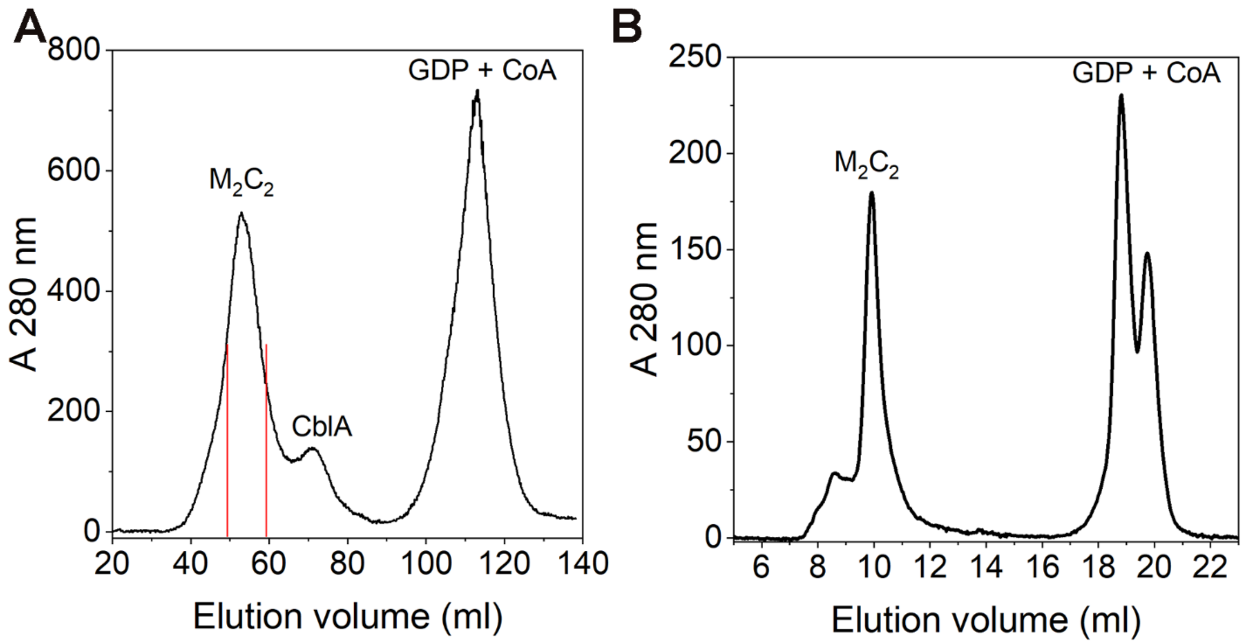


**Figure S9. Purification of the human $M_2C_2$ complex for crystallization.** (**A**) Gel filtration profile of 100 μM MMUT, 200 μM MMAA, 2 mM GDP and 2 mM CoA on an S300 column. The $M_2C_2$ peak fractions (red bars) were used for crystallization. **(B).** The purified $M_2C_2$ complex from **A** was reanalyzed on an analytical S200 column, which demonstrated that the complex was intact.
